# JAK2/STAT3 signaling pathway mediates methylmercury toxicity in mouse astrocyte neuronal C8-D1A cell line

**DOI:** 10.1101/2024.07.13.603400

**Authors:** Aafia Ahmed, Michael Aschner, Beatriz Ferrer

## Abstract

Methylmercury (MeHg) is an environmental pollutant. Consumption of contaminated fish is the main exposure route in humans, leading to severe neurological disorders. Upon ingestion MeHg reaches the brain and selectively accumulates in astrocytes disrupting glutamate and calcium homeostasis and increasing oxidative stress. Despite extensive research, the molecular mechanisms underlying MeHg neurotoxicity remain incompletely understood. The induction of nuclear factor erythroid 2-related factor 2 (Nrf2) and its role activating antioxidant responses during MeHg-induced oxidative injury have garnered significant attention as a potential therapeutic target against MeHg toxicity. However, recent studies indicate that the Nrf2 signaling pathway alone may not be sufficient to mitigate MeHg-induced damage, suggesting the existence of other protective mechanisms. The signal transducer and activator of transcription 3 (STAT3) plays a crucial role in cell growth and survival. Several studies have also highlighted its involvement in regulating redox homeostasis, thereby preventing oxidative stress through mechanisms that involve modulation of nuclear genes that encode electron transport complexes (ETC) and antioxidant enzymes. These characteristics suggest that STAT3 could serve as a viable mechanism to mitigate MeHg toxicity, either in conjunction with or as an alternative to Nrf2 signaling. Our previous findings demonstrated that MeHg activates the STAT3 signaling pathway in the GT1-7 hypothalamic neuronal cell line, suggesting its potential role in promoting neuroprotection. Here, to elucidate the role of the STAT3 signaling pathway in MeHg neurotoxicity, we pharmacologically inhibited STAT3 using AG490 in the C8D1A astrocytic cell line exposed to 10 µM MeHg. Our data demonstrated that pharmacological inhibition of STAT3 phosphorylation exacerbates MeHg-induced mortality, antioxidant responses, and ROS production, suggesting that STAT3 may contribute to neuroprotection against MeHg exposure in astrocytes.

## 1. Introduction

Minamata disease, first discovered in the city of Minamata, Japan, is a neurological disease resulting from high mercury consumption (Eto, Marumoto et al. 2010). In the late 1950s, large amounts of water were tainted with mercury due to waste from a nearby chemical plant owned by the Japanese corporation Chisso, resulting in the bioaccumulation and biomagnification of methylmercury in the muscles of fish (Harada 1995, McCurry 2006). Those impacted by methylmercury contamination reported the sudden onset of symptoms like sensory disturbances, ataxia, dysarthria, and tunnel vision (Harada 1995, Clarkson, Magos et al. 2003, Grandjean, Satoh et al. 2010). These symptoms generally worsened and were followed by seizures, a coma, and ultimately, death (Bakir, Rustam et al. 1980, Yuan 2012). By the end of 1980, 378 of the 1422 Minamata disease patients had passed away (Tamashiro, Akagi et al. 1984). This event revealed the toxic nature of mercury and emphasized the importance of effectively controlling its levels and averting future instances of mercury poisoning.

Mercury is naturally found within Earth’s crust (World Health Organization 2017). For certain species like wild piscivorous fish, mammals, and birds, toxicological impacts include altered growth, reproduction, neurodevelopment, learning capacity, and behavioral abnormalities that raise mortality and predation risks (Wren 1986, Depew, Burgess et al. 2013). Inorganic mercury is deposited on land by attaching to airborne particles in various forms of precipitation. Mercury is then returned into the atmosphere in a different form and redeposited. During this ongoing cycle, mercury undergoes complex chemical and physical changes. Microscopic organisms in water bodies combine mercury with carbon to form organic mercury. The most common type of organic mercury, methylmercury, is found in the environment and is highly toxic (Environmental Protection Agency 2023).

Methylmercury (MeHg) can directly interact with brain and fetal cells by crossing the blood-brain and placental barriers (Cariccio, Sama et al. 2019). In the gastrointestinal tract, MeHg forms a complex with L-Cysteine (CH_3_Hg-Cys) by imitating the amino acid methionine, enabling it to simply pass through the blood-brain barrier (Kerper, Ballatori et al. 1992). MeHg has a high affinity to sulfhydryl (-SH) protein groups, making it difficult to eliminate MeHg from cells (Ajsuvakova, Tinkov et al. 2020). The kidneys and nervous systems are particularly susceptible to methylmercury poisoning (Fernandes Azevedo, Barros Furieri et al. 2012).

The key mechanisms involved in MeHg-induced toxicity include increased reactive oxygen species, biomarkers of oxidative stress, and superoxide dismutase (SOD) activities. The activation of antioxidant molecules may defend the central nervous system from these neurotoxic symptoms of MeHg (do Nascimento, Oliveira et al. 2008, Liao, Peng et al. 2021). Previously, upregulation of the nuclear factor erythroid 2-related factor 2 (Nrf2) and the downstream genes have been implicated in MeHg toxicity (Ni, Li et al. 2011). However, many studies demonstrate the involvement of different redox signaling in MeHg-induced neurotoxicity, including the mitogen-activated protein kinase (MAPK) cascade and Rho-associated coiled coil-forming protein kinase (ROCK) signaling (Fujimura, Usuki et al. 2019, Takanezawa, Sakai et al. 2023), suggesting the involvement of multiple pathways to combat MeHg toxicity.

The seven members (STAT1, STAT2, STAT3, STAT4, STAT5a, STAT5b, and STAT6) of the signal transducer and activator of transcription (STAT) proteins serve as innovative therapeutic targets for the treatment of diseases (Wang, Man et al. 2022, Wong, Manore et al. 2022). STAT-3 is crucial for cell cycle, cell proliferation, cell apoptosis, and the development of tumors (Gu, Mohammad et al. 2020). It is activated by phosphorylation and translocated to the nucleus to participate in the regulation of gene expression. STAT-3 can be regulated by upstream signaling molecules, such as Janus kinase (JAK) and epidermal growth factor receptor (EGFR). It then localizes to the nucleus of cells and binds to target DNA to regulate the expression of downstream proteins (Wang, Man et al. 2022). STAT-3 acts as a downstream response protein for multiple cytokines and modulates cellular proliferation and intercellular interactions (Li, Yin et al. 2022). Due to its critical role in a variety of biological processes, including angiogenesis, differentiation, anti-apoptosis, proliferation, and inflammatory response, this signal transducer is the subject of extensive research.

However, the STAT-3 signaling pathway has been poorly studied in the context of MeHg toxicity in an astrocytic model. Low concentrations of MeHg enhance ciliary neurotrophic factor (CNTF)-evoked STAT3 phosphorylation in human neuroblastoma SH-SY5Y and mouse cortical neural progenitor cells (NPCs) (Jebbett, Hamilton et al. 2013). Additionally, persistent MeHg exposure via drinking water triggers STAT3 phosphorylation in the hypothalamus of mice (Ferrer, Prince et al. 2021), and prolonged exposure to MeHg-albumin conjugates increases STAT3 phosphorylation in microglial N9 cells (Tan, Zhang et al. 2019). Notably, the hypothalamic neuronal cell line GT1-7 provided the first evidence of a mechanistic correlation between MeHg-induced oxidative stress and STAT3 signaling (Ferrer, Suresh et al. 2021). Interestingly, MeHg has also been associated with amyotrophic lateral sclerosis (ALS)-like disease (Carocci, Rovito et al. 2014, Minj, Upadhayay et al. 2021), which involve elevated STAT-3 protein levels in the brain in rats (Kumar, Mehan et al. 2024). Despite these findings, the precise role of STAT-3 activation in MeHg toxicity specifically within astrocytes remains unclear.

Despite its established role in various cellular processes, the involvement of the STAT-3 signaling pathway in astrocyte-mediated MeHg neurotoxicity remains elusive. Although studies have documented MeHg-induced STAT3 phosphorylation in diverse cell types, including NPCs, hypothalamic neurons, and microglia (Jebbett, Hamilton et al. 2013, Tan, Zhang et al. 2019, Ferrer, Suresh et al. 2021), a definitive link between STAT-3 activation and MeHg-mediated astrocytic damage is lacking. This knowledge gap is particularly concerning given the crucial role astrocytes play in maintaining central nervous system (CNS) homeostasis and their established susceptibility to MeHg exposure. To address the specific role of STAT-3 in the astrocytic response to MeHg, we employed an in vitro approach using C8-D1A astrocyte cells. Tyrphostin B 42 (AG490; 2-cyano-3-(3,4-dihydroxyphenyl)-N-(phenylmethyl)-2-propenamide), a selective inhibitor of the JAK2 tyrosine kinase (Meydan, Grunberger et al. 1996), known to decrease STAT3 phosphorylation, will be used to dissect the STAT-3 pathway’s contribution. We hypothesized that MeHg will induce STAT3 phosphorylation, potentially playing an antioxidant role in astrocytes. Conversely, disruption of STAT3 signaling through JAK2 inhibition with AG490 may exacerbate MeHg toxicity by increasing reactive oxygen species (ROS) production and cellular death. Astrocytic C8-D1A cells were exposed to MeHg (0 or 10µM) and AG490 (0, 10, 50, or 100 µM) at varying exposure times.

## 2. Material and methods

### 2.1. Cell culture

Mouse astrocytic neuronal C8-D1A cell lines were obtained from the American Type Cell Culture Collection (ATCC, CRL-2541) and grown in Dulbecco’s modified Eagles’ medium (DMEM) (Gibco™, 11995040) supplemented with 10% heat-inactivated fetal bovine serum (Gibco™, 10438034) and 1% penicillin/streptomycin (Gibco™, 15140148). Cells were maintained at 37 °C with 5% CO_2_.according to the ATCC mammalian tissue culture protocol and sterile techniques.

### 2.2. Methylmercury (MeHg) exposure

Cells were subcultured and treated with methylmercury (II) chloride (MeHg) (Sigma-Aldrich, 442534) at 0 (control) or 10µM for 1, 3, and 24h unless otherwise specified.

### 2.3. STAT3 inhibitor treatment

Cells were pre-treated for 1h with STAT3 inhibitor AG490 (Tyrphostin B42 or 2-cyano-3-(3,4-dihydroxyphenyl)-N-(phenylmethyl)-2-propenamideor), Santa Cruz Biotechnology, sc-202046) at 0, 10, 50, or 100 µM, followed by co-treatment with 0 or 10µM MeHg at specified times.

### 2.4. Lactate dehydrogenase (LDH) cytotoxicity assay

C8-D1A cells were plated in a 96-well plate. Lactate dehydrogenase (LDH) released from the cell was used to measure MeHg cytotoxicity. LDH released from the cell oxidizes lactate to generate NADH, which reacts with iodonitrotetrazolium (INT) to form the red-colored formazan salt (Smith, Wunder et al. 2011). 24h after seeding, cells were treated with assigned concentrations of MeHg and AG490. LDH release was measured in the medium using the CyQUANT LDH Cytotoxicity Assay Kit. 50 µL of each well with AG490 and MeHg was transferred to a new plate and 50 µL of LDH reaction mixture was prepared, with an aliquot added to the new plate. The plate was incubated for 30 minutes under a fume hood protected from light, followed by the addition of 50 µL of stop solution. Absorbance was measured at 490nm and 680nm using a microplate reader.

### 2.5. 3-(4,5-dimethylthiazol-2-yl)-2,5-diphenyltetrazolium bromide (MTT) cell viability assay

Cell viability was measured with 3-(4,5-dimethylthiazol-2-yl)-2,5-diphenyltetrazolium bromide (MTT) (Thermo Fisher Scientific, M6394). After cell exposure to AG490 and MeHg at specified times and concentrations, 100 µL of 0.5 µg/µL MTT in Hank’s balanced salt solution (HBSS) (Gibco™, 14175095) for one hour and 30 minutes at 37LJ with 5% CO_2_. Then the supernatant was removed and MTT was solubilized in dimethyl sulfoxide (DMSO) (Sigma Aldrich, D8418). Absorbance was measured at 540 nm using a microplate reader.

### 2.6. Reactive oxygen species (ROS) production

Cellular ROS production was assessed using a general oxidative stress indicator CM-H_2_DCFDA (Thermo Fisher Scientific, C6827). Cells were seeded in a black 96-well microplate and labeled with 5 µM CM-H_2_DCFDA diluted in HBSS followed by treatment with STAT3 inhibitor and MeHg. Fluorescence was measured at 0h, 30min, 1h, 3h, 6h, and 24h after MeHg addition at excitation 495 nm and emission 525 nm using a microplate reader.

Mitochondrial ROS production was visualized using MitoSOX (Thermo Fisher Scientific, M36008). Cells were seeded in a black 96-well microplate and labeled with 500 nM MitoSOX diluted in HBSS. Cells were treated with STAT3 inhibitor for 1h, followed by exposure to 10 µM MeHg. Fluorescence was measured at 0h, 30 min, 1h, 3h, and 6h after addition of MeHg using a microplate reader.

### 2.7. Western blot

Cells were sub-cultured at 9 x 10^5^ cells into 60 mm culture dish. For whole cell protein extracts, cold radioimmunoprecipitation assay (RIPA) buffer (Sigma Aldrich, R0278) was added with phosphatase inhibitor cocktails 2 and 3 (Sigma Aldrich, P5726 and P0044, respectively) and Halt™ protease inhibitor cocktail (Thermo Fisher Scientific, 78437) was used. The lysates were sonicated and centrifuged. The supernatant was collected, and protein concentration was measured using the Pierce BCA Protein Assay Kit (Thermo Fisher Scientific, 23225). Samples were dissolved in Laemmli sample buffer (Bio-Rad, #1610737) and boiled at 100℃ for 5 minutes, 20 µg of protein was resolved by 4-20% sodium dodecyl sulfate-polyacrylamide gel (SDS-PAGE, Bio-Rad #4561096). The gels were then transferred to a nitrocellulose membrane and incubated for one hour with 5% bovine serum albumin (BSA, Fisher Scientific, BP1600) in Tris buffer saline-0.1% tween-20 (TBS-T) at room temperature to block non-specific binding, followed by incubation with primary antibodies diluted in 5% BSA in TBS-T buffer overnight at 4℃. The blots were washed three times and incubated with selected horseradish peroxidase-conjugated secondary antibody. Protein bands were detected with a chemiluminescent method (Thermo Fisher Scientific 34579 or 37071). ImageJ software (NIH) was used to analyze the bands. For whole lysates, densitometry was normalized to β-actin as a housekeeping protein. Antibodies used included: mouse anti-β-actin (Sigma Aldrich, A1978), rabbit anti-phospho-STAT3 (Tyr705) (Cell Signaling Technology, #9145), mouse anti-STAT3 (Cell Signaling Technology, #9139), anti-Nrf2 (Cell Signaling Technology, #12721), and mouse anti-HO1 (Cell Signaling Technology, #86806).

### 2.8. Statistical analysis

Statistical analyses were performed using GraphPad Prism software and/or Excel. Statistical analysis was completed using two-way ANOVA, followed by Tukey’s *post-hoc* test. All data are reported as mean ± standard error (SD) and statistical significance was defined as p < 0.05.

## 3. Results

### 3.1. Inhibition of STAT3 phosphorylation exacerbates MeHg-induced death in C8-D1A astrocytic cells

AG490 selectively blocks JAK2, inhibiting the activity of STAT3. The impact of STAT3 inhibition on cell survival, death, and ROS production was examined to elucidate the role of STAT3 in MeHg-induced toxicity. C8D1A astrocytic cells were pre-treated for 1h with 0, 10, 50, or 100 µM AG490, followed by exposure to 10 µM MeHg. Cytotoxicity was measured using LDH assay at 3, 6, and 24h (Figure 1A, 1B, and 1C, respectively). No significant effects were detected at 3h or 6h. However, at 24h MeHg exposure significantly increased cytotoxicity in C8D1A astrocytic cells across all concentrations of the inhibitor (F_(1,40)_= 33.307, p < 0.001) (Figure 1C).

**Figure 1.**
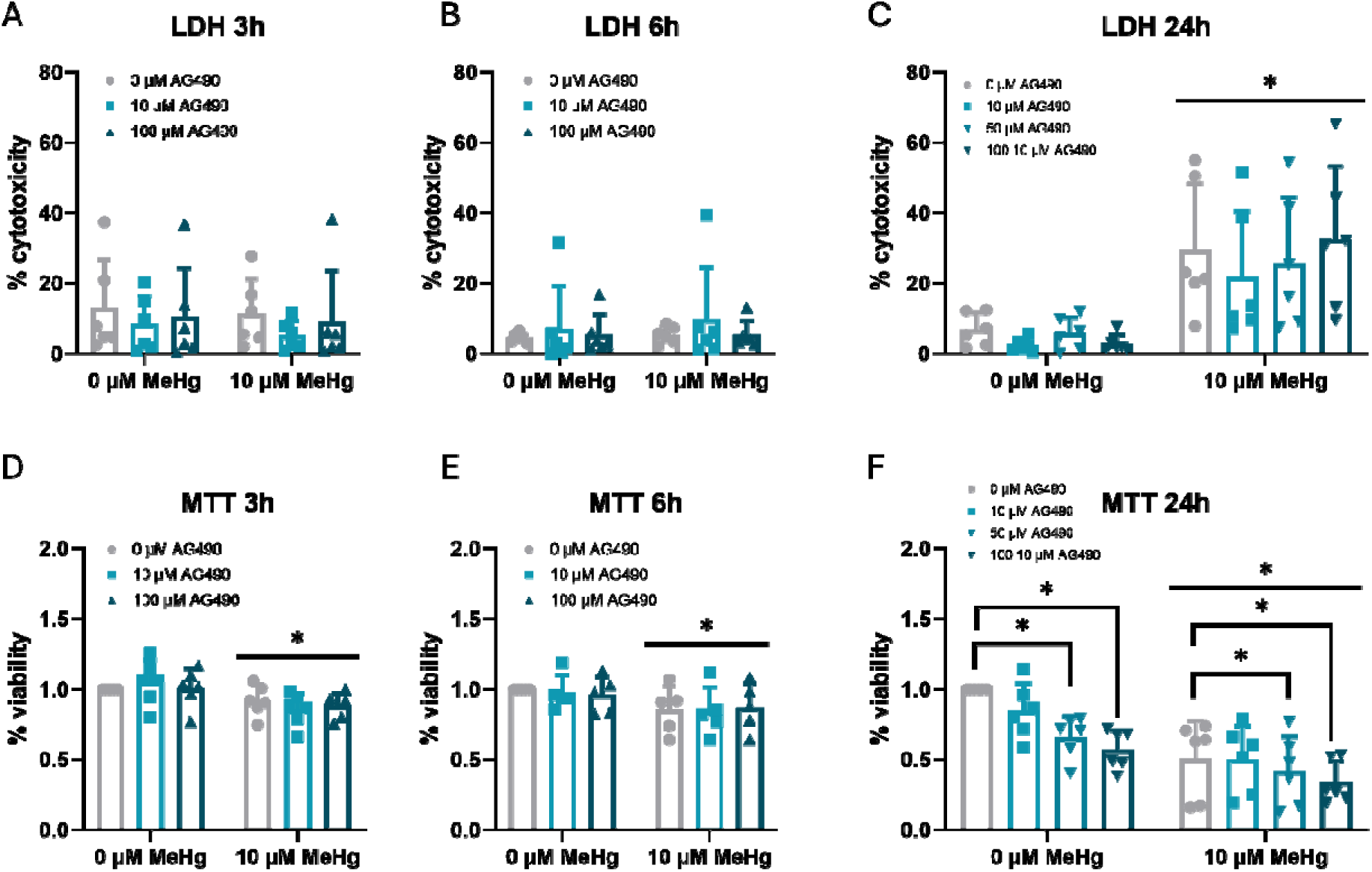
Inhibition of STAT3 phosphorylation enhanced MeHg toxicity in C8-D1A cells. Astrocytic C8-D1A cells were treated with AG490 (0, 10, 50, or 100µM) for one hour followed by co-exposure to MeHg (0 or 10µM) for 3 hours (A and D), 6 hours (B and E), and 24 hours (C and F). Cytotoxicity was measured using the LDH assay (A, B, and C), and viability was evaluated using the MTT assay (D, E, and F). Data are represented as mean ± SD. Data were analyzed by two-way ANOVA followed by Tukey’s *post hoc* test. * denotes statistically significant compared to the respective control (p < 0.05).

Cell viability was assessed using a MTT assay at 3, 6, and 24h (Figure 1D, 1E, and 1F, respectively). MeHg exposure significantly decreased cell viability at 3h (F_(1,30)_= 12.340, p = 0.001), 6h (F_(1,25)_= 6.329, p = 0.019), and 24h (F_(1,40)_= 35.563, p < 0.001) (Figure 1D, 1E, and 1F). Cell viability steadily decreased when inhibitor concentrations were increased in cells not exposed to MeHg, indicating that the JAK2-specific inhibitor may generally increase susceptibility to cellular damage. Additionally, in MeHg-exposed cells, increasing concentrations of inhibitor further exacerbated cell death (F_(1,40)_= 5.979, p = 0.002) (Figure 1F). This suggests a possible interaction between AG490 and MeHg, leading to enhanced cytotoxicity.

LDH assay measures lactate dehydrogenase (LDH) release, an indicator of compromised membrane integrity. Conversely, the MTT assay assesses metabolic activity within the mitochondria. Differences observed between LDH and MTT results suggest that STAT3 inhibition may exacerbate MeHg toxicity through mitochondrial dysfunction.

### 3.2. Inhibition of STAT3 phosphorylation increases MeHg-induced ROS production in C8-D1A astrocytic cells

The effect of STAT3 inhibition on MeHg-induced intracellular ROS production was also measured (Figure 2 and 3). First, we measured overall cellular ROS production using CM-H2DCFDA. Consistent with our expectations, MeHg exposure resulted in a significant elevation of ROS levels at 3h (F_(1,64)_ = 13.040, p < 0.001) (Figure 2D), persisting at 6h (F_(1,46)_ = 5.640, p = 0.022) (Figure 2E), and 24h (F_(1,32)_ = 18.450, p < 0.001). Interestingly, inhibitor treatment exacerbated MeHg-induced ROS production at 24h (F_3,32)_ = 3.987, p = 0.016) (Figure 2F).

**Figure 2.**
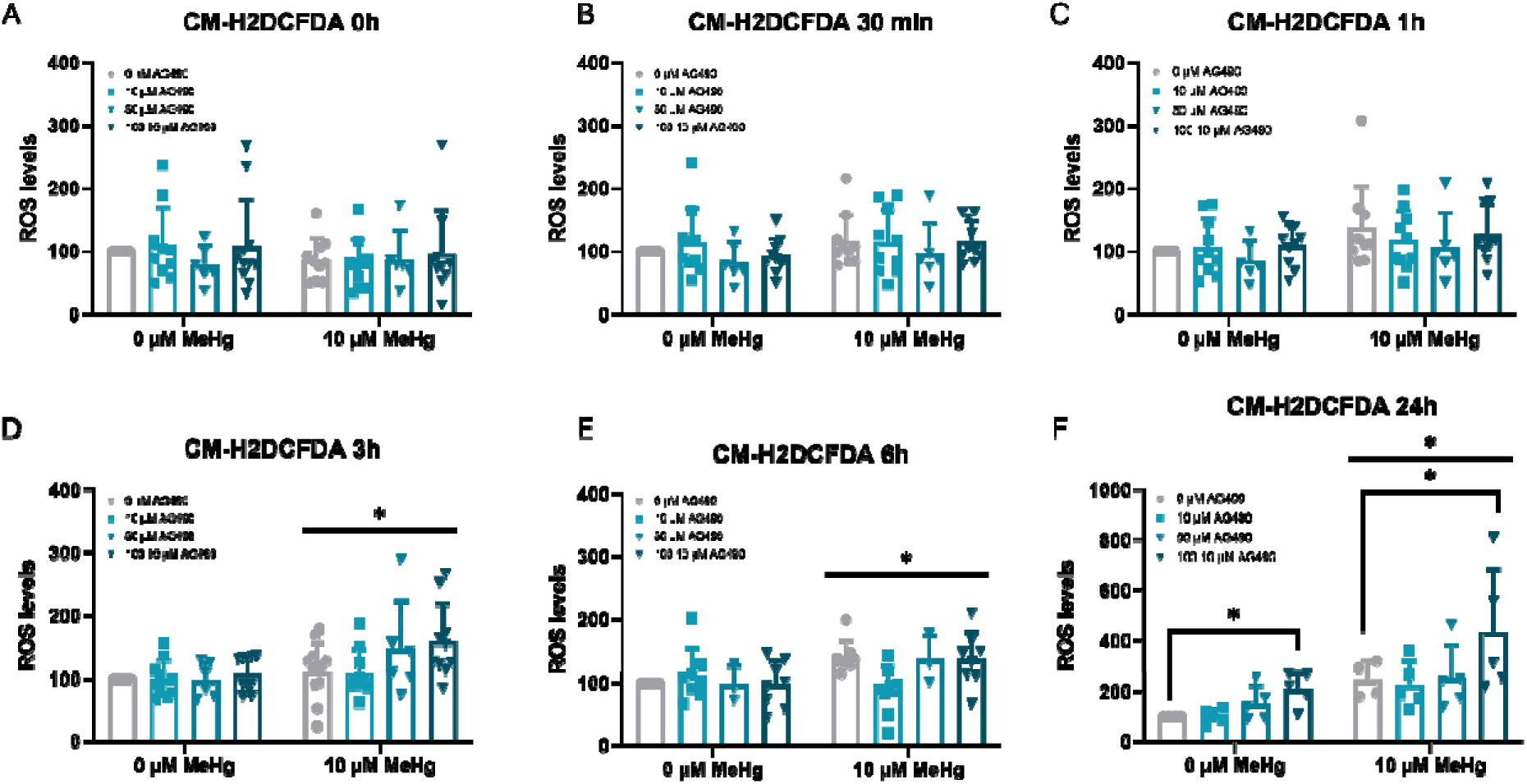
Blocking STAT3 phosphorylation enhances MeHg-induced intracellular ROS production in C8-D1A cells. Astrocytic C8-D1A cells were treated with AG490 (0, 10, 50, or 100µM) for one hour followed by co-exposure to MeHg (0 or 10µM) for 0 hours (A), 30 min (B), 1 hours (C), 3 hours (D), 6 hours (E), and 24 hours (F). Intracellular ROS was measured using the CM-H_2_DCFDA fluorescent probe. Data are represented as percentage of control as mean ± SD. Data were analyzed by two-way ANOVA followed by Tukey’s *post hoc* test. * denotes statistically significant compared to the respective control (p < 0.05).

**Figure 3.**
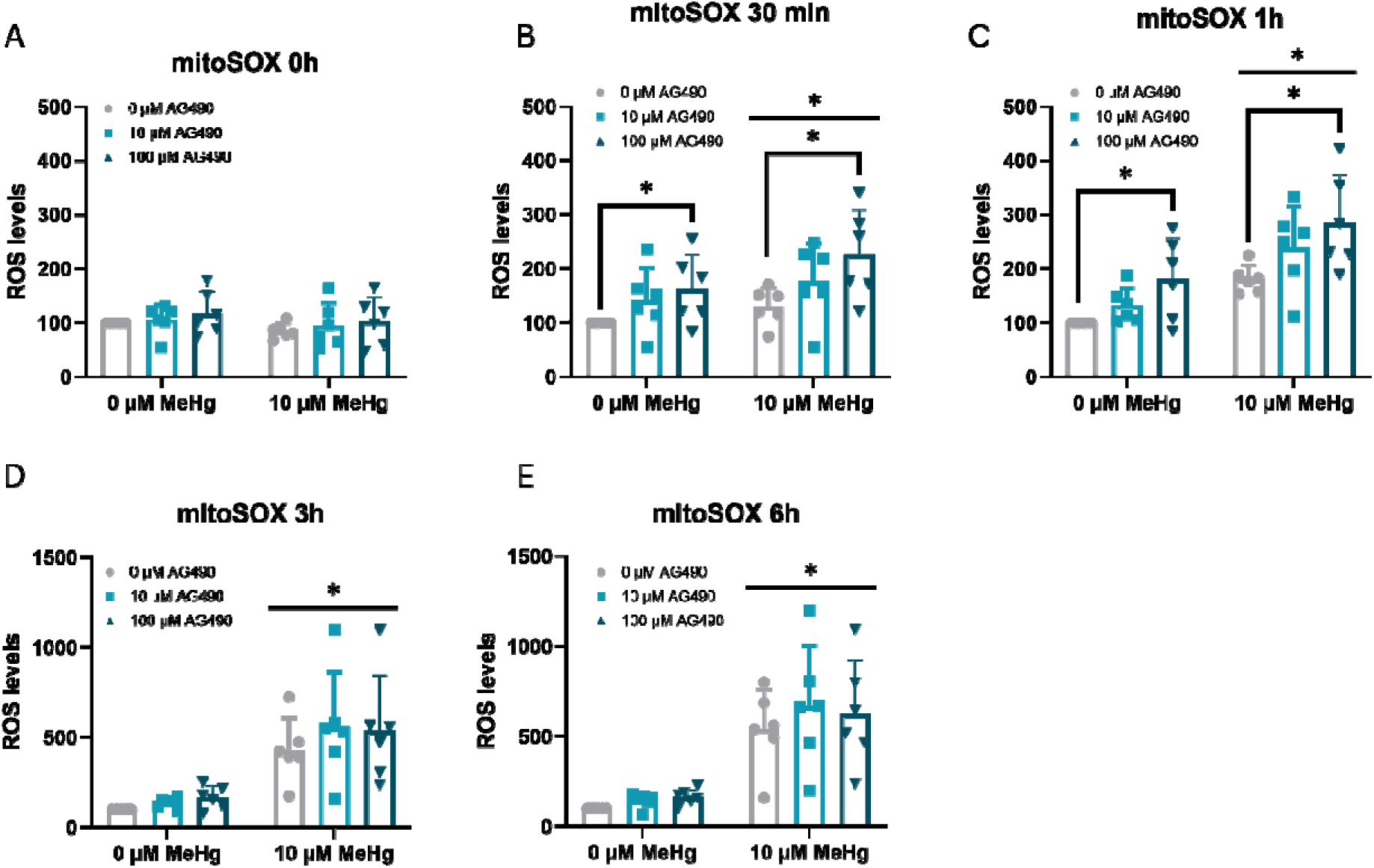
Inhibition of STAT3 phosphorylation aggravates MeHg-induced mitochondrial ROS production in C8-D1A cells. Astrocytic C8-D1A cells were treated with AG490 (0, 10, 50, or 100µM) for one hour followed by co-exposure to MeHg (0 or 10µM) for 0 hours (A), 30 min (B), 1 hours (C), 3 hours (D), and 6 hours (E). Mitochondrial ROS was measured using the mitoSOX fluorescent probe. Data are represented as percentage of control as mean ± SD. Data were analyzed by two-way ANOVA followed by Tukey’s *post hoc* test. * denotes statistically significant compared to the respective control (p < 0.05).

Next, we assessed mitochondrial oxidative stress specifically using MitoSOX. Our data revealed that MeHg significantly increased mitochondrial superoxide levels at 30 min (F_(1,30)_ = 5.057, p = 0.032) (Figure 3B), with the effect persisting at 1h (F_(1,30)_ = 24.176, p < 0.001) (Figure 3C), 3h (F_(1,30)_ = 34.040, p < 0.001) (Figure 3D), and 6h (F_(1,30)_ = 48.792, p < 0.001) (Figure 3F). Notably, the AG490 inhibitor increased superoxide production at 30 min (F_2,30)_ = 5.460, p = 0.009) (Figure 3B) and at 1h (F_(2,30)_ = 17.315, p = 0.003) (Figure 3C).

These findings suggest that altered STAT3 signaling contributes to elevated ROS production, potentially in part through disruption of mitochondrial homeostasis. This observation strengthens the hypothesis that STAT3 plays a key role in maintaining cellular redox balance, and its alterations result in oxidative stress and irreversible cellular damage.

### 3.3. STAT3 phosphorylation and Heme Oxygenae-1 (HO-1) induction upon MeHg exposure and STAT3 inhibition in C8-D1A astrocytic cells

To fully elucidate the role of STAT3 in cellular defense against oxidative stress, cells were pretreated with 0, 10, 50, or 100 µM AG490 for one hour followed by co-treatment with 0 or 10 µM MeHg for one hour. Protein expression was then assessed using immunoblot. First, phosphorylated and total STAT3 protein levels were assessed. Phospho-STAT3 expression significantly increased in all cells treated to 10 µM MeHg (F_(1,56)_ = 106.645, p < 0.001) (Figure 4A). Surprisingly, no significant differences were observed in phospho-STAT3 levels between MeHg-exposed cells and those co-treated with increasing concentrations of the inhibitor (AG490). As expected, total STAT3 protein expression remained unaffected by either MeHg or the AG490 inhibitor treatment (Figure 4B). Following 1 hour of MeHg exposure, the ratio of phospho-STAT3 to total STAT3 significantly increased (F_(1,56)_ = 190.833, p < 0.001) (Figure 4C), indicating MeHg-induced STAT3 activation. Importantly, AG490 treatment failed to inhibit MeHg-mediated STAT3 phosphorylation.

**Figure 4.**
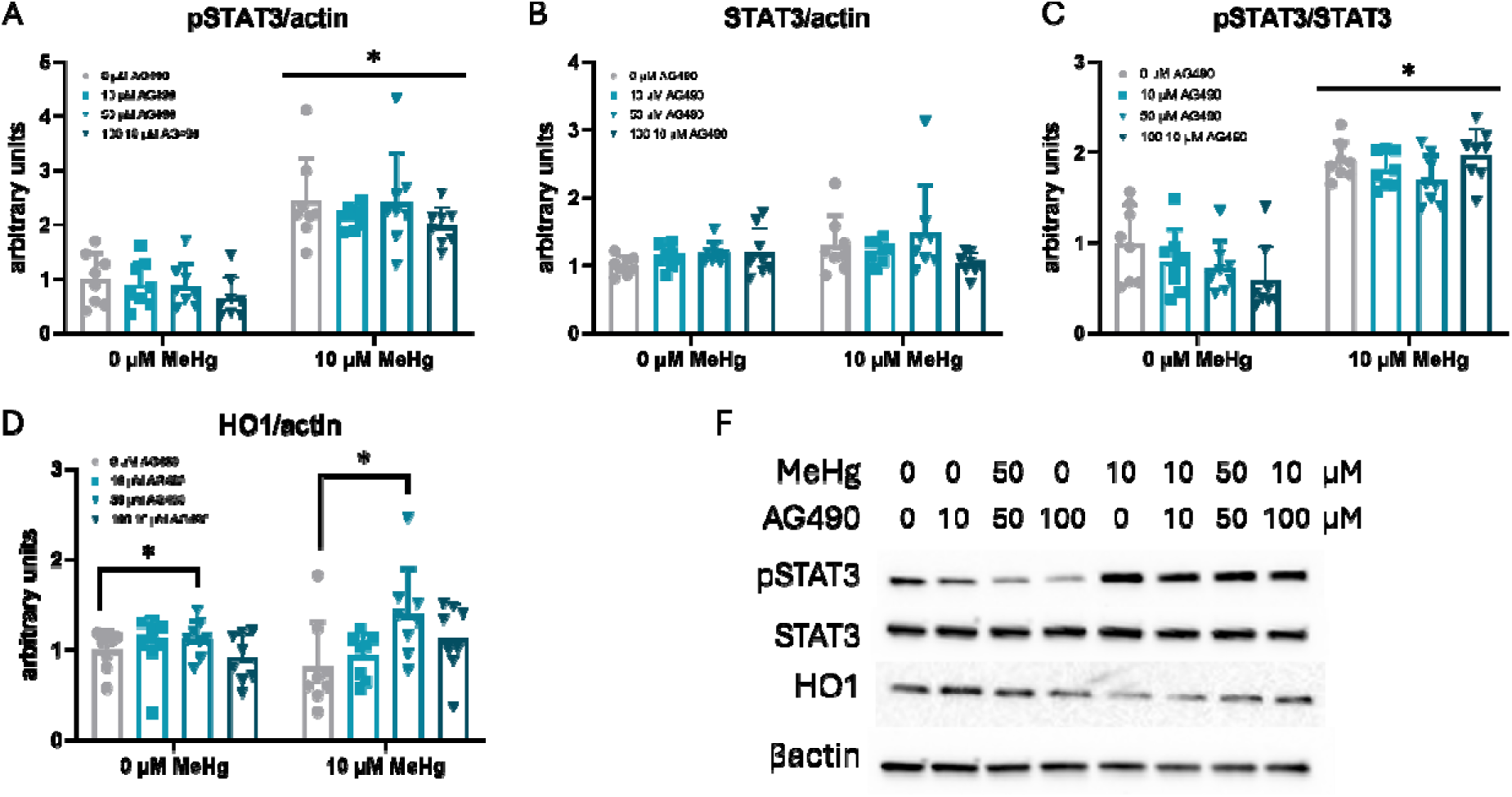
Inhibition of STAT3 phosphorylation exacerbates MeHg-induced HO1 protein expression in C8-D1A cells. Astrocytic C8-D1A cells were treated with AG490 (0, 10, 50, or 100µM) for one hour followed by co-exposure to MeHg (0 or 10µM) for 1 hour. Western blot analysis was performed to measure (A), pSTAT3, (B), STAT3, (C) pSTAT3/STAT3 ratio, and (D) HO1 protein expression levels, quantified by densitometry. (E), Representative western blot of the total and phosphorylated proteins are shown. Data are presented as percentage of control as mean ± SD. Data were analyzed by two-way ANOVA followed by Tukey’s *post hoc* test. * denotes statistically significant compared to the respective control (p < 0.05).

Heme oxygenase-1 (HO-1), a key antioxidant enzyme regulated by the Nrf2 pathway, is responsible for heme group degradation (Funes, Rios et al. 2020). MeHg exposure for 1h did not significantly affect HO-1 protein levels. However, AG490 inhibitor significantly increased HO-1 expression in a concentration-dependent manner (F_(3,56)_ = 2.980, p = 0.039) (Figure 4D), suggesting a direct link between activation of the JAK2 pathway and the Nrf2 signaling pathway.

We further investigated STAT3 phosphorylation and HO-1 induction following a longer exposure time (3h). Cells were pretreated with increasing concentrations of AG490 (0, 10, or 100 µM) for 1h, followed by co-treatment with 0 or 10 µM MeHg for 3h. Similar to the one-hour exposure, MeHg exposure for 3h significantly induced both STAT3 phosphorylation (F_(1,24)_ = 46.897, p < 0.001) (Figure 5A) and phospho-STAT3/STAT3 ratio (F_(1,24)_ = 50.790, p < 0.001) (Figure 5C), indicating sustained STAT3 activation by MeHg. However, consistent with the one-hour data, AG490 inhibitor again failed to inhibit MeHg-mediated STAT3 phosphorylation (Figure 5A and 5CD). As expected, total STAT3 protein levels remained unaffected by MeHg and the JAK2 inhibitor (Figure 5B).

**Figure 5.**
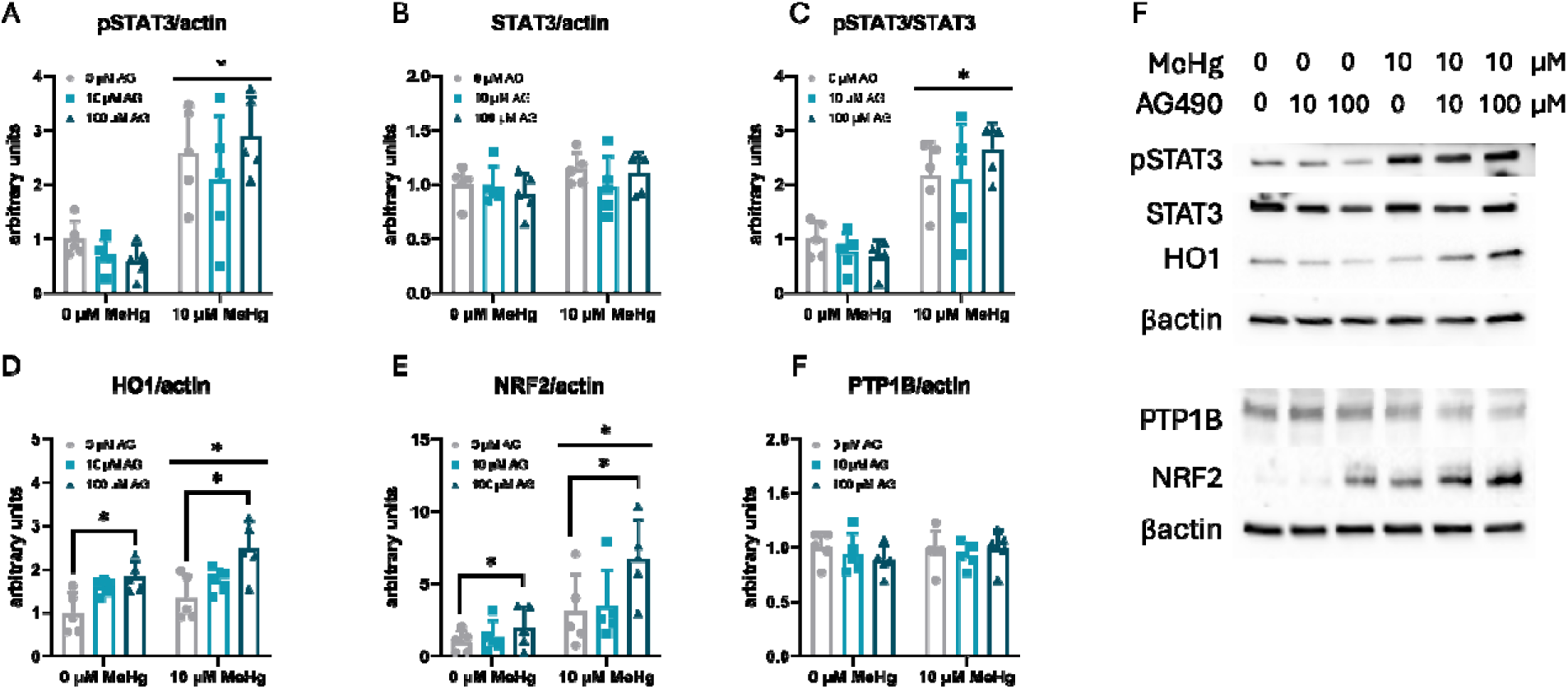
Inhibition of STAT3 phosphorylation exacerbates MeHg-induced HO1 protein expression in C8-D1A cells. Astrocytic C8-D1A cells were treated with AG490 (0, 10, 50, or 100µM) for one hour followed by co-exposure to MeHg (0 or 10µM) for 3 hours. Western blot analysis was performed to measure (A), pSTAT3, (B), STAT3, (C) pSTAT3/STAT3 ratio, (D) HO1, (F) Nrf2 protein expression levels, quantified by densitometry. (F), Representative western blot of the total and phosphorylated proteins are shown. Data are presented as percentage of control as mean ± SD. Data were analyzed by two-way ANOVA followed by Tukey’s *post hoc* test. * denotes statistically significant compared to the respective control (p < 0.05).

Interestingly, HO-1 protein levels were significantly increased after 3h of MeHg exposure (F_(1,24)_ = 5.550, p = 0.027) (Figure 5D). Furthermore, the higher dose of AG490 (100 µM) further enhanced this MeHg-induced HO-1 upregulation (F_(2,24)_ = 13.013, p < 0.001) (Figure 5D). To gain insights into the mechanism by which AG490 mediates HO-1 activation, we quantified Nrf2 protein levels, the transcription factor responsible for HO-1 gene expression. MeHg exposure increased Nrf2 protein expression (F_(1,24)_ = 16.858, p < 0.001) (Figure 5E), and this increase was further augmented by AG490 treatment (F_(2,24)_ = 3.924, p = 0.034) (Figure 5E).

Since AG490 targets the JAK2 pathway, we also examined protein tyrosine phosphatase 1B (PTP1B), a known JAK2 inhibitor (Myers, Andersen et al. 2001). Neither MeHg nor AG490 treatment affected PTP1B expression (Figure 5F).

Altogether these results suggest that JAK2 inhibition potentially mediates Nrf2-HO-1 signaling pathway through a mechanism independent of STAT3 activation.

## 4. Discussion

The signaling pathways involved in the deleterious effect of MeHg toxicity remain unclear. This study demonstrates the antioxidant role of the STAT3 signaling pathway for oxidative stress regulation of MeHg-induced toxicity. MeHg has been revealed in numerous studies to modify protein phosphorylation in a concentration- and cell-type-dependent manner, resulting in the dysregulation of signaling pathways (Sarafian and Verity 1990, Yagame, Horigome et al. 1994, Ke, Goncalves et al. 2019). However, very little is known about the effect of MeHg on the STAT3 signaling pathway. Tan et al found that ERK/MAPKs and STAT3 signaling pathways may be related to the hormesis of MeHg-HSA in N9 cells (Tan, Zhang et al. 2019), while Jebbett and colleagues found that specific levels of MeHg enhanced CNTF-evoked STAT3 phosphorylation, target-gene expression, and glial differentiation (Jebbett, Hamilton et al. 2013). For the first time, the role of STAT3 in MeHg toxicity was implicated in hypothalamic neuronal cells where MeHg induced STAT3 phosphorylation in a time- and concentration-dependent manner (Ferrer, Suresh et al. 2021). However, STAT3 activation has yet to be demonstrated in other cell models. Therefore, this study aimed to investigate the role of STAT3 on MeHg-induced toxicity in an astrocytic cell model.

Our study investigated the role of STAT3 signaling in MeHg-induced oxidative stress. We found that inhibiting STAT3 pharmacologically led to a significant increase in ROS production within MeHg-exposed cells. These findings align with a previous study indicating that the disruption of astrocyte STAT3 signaling decreases mitochondrial function and increases oxidative stress *in vitro* (Sarafian, Montes et al. 2010). Thus, altering the STAT3 signaling pathway results in increased ROS production. During oxidative stress conditions, Nrf2 translocates into the cell nucleus, forms a dimer with sMaf, and binds to antioxidant response elements (AREs) located in the regulatory regions of genes responsible for cellular defense (Unoki, Akiyama et al. 2018). Numerous studies have revealed an indirect relationship between STAT3 and Nrf2, wherein decreased STAT3 phosphorylation results in increased Nrf2 expression (Mohamed, Hafez et al. 2018). The relationship between STAT3 and Nrf2 is also supported by computational analysis of the connection between Nrf2 and STAT1 and STAT3 (Turei, Papp et al. 2013). Additionally, Wruck et al. (2011) propose an interaction between Nrf2 and IL6, suggesting that Nrf2 may play a role in inflammatory responses by binding to antioxidant response elements in the promoter region of the IL6 gene and activating its transcription (Wruck, Streetz et al. 2011). Taken together, findings suggest a link between antioxidant and anti-inflammatory mechanisms. In this study, AG490 inhibitor enhanced the MeHg-induced expression of HO-1 supports the notion that Nrf2 is activated when JAK2 pathway is inhibited. Further research is needed to investigate the implication of anti-inflammatory genes such as the *Il6* gene in the defense response against MeHg toxicity.

As demonstrated in this study, inhibition of STAT3 phosphorylation resulted in the decrease of cell viability in a concentration-dependent manner. STAT3 has been shown to play a neuroprotective role against glutamate-induced excitotoxicity in primary cultures of dorsal root ganglion neurons and glioblastoma cells (Wen, Li et al. 2017, Corbetta, Di Ianni et al. 2019). Activated STAT3 promotes cell survival and proliferation through the upregulation of Cyclin D1, Pim1, and Bcl2 (Lu, Dong et al. 2018). Further research is warranted to evaluate how gene expression is affected by MeHg. Previously, it had been revealed that the increase in MeHg mortality is correlated with the decrease of *Ccnd1* and *Pim1* mRNA after inhibition of JAK2/STAT3 (Ferrer, Suresh et al. 2021).

As expected, HO-1 levels were also increased in MeHg-exposed cells compared to the control, suggesting the activation of the N2 signaling pathway. This is consistent with the finding that NRF2 is a regulator of cellular resistance and acts as a defense against toxicity through pharmacological interventions (Ma 2013). Previously, activation of the Nrf2/HO-1 signaling pathway, inhibited excessive oxidative stress and inflammation (Yang, Yin et al. 2019, Minj, Upadhayay et al. 2021). This study demonstrated that regulation of Nrf2/HO-1 activation may allow for the development of therapeutic drugs to treat joint conditions.

In conclusion, the data presented in this study demonstrate the neuroprotective role of STAT3 as a defense response in MeHg-induced toxicity. Increasing concentrations of inhibitor increased ROS production levels in MeHg exposed C8-D1A cells, decreased cell proliferation, and increased cytotoxicity.

Additionally, there was an increase in HO-1, suggesting their activation as a defense response in MeHg-toxicity. Future research is warranted to fully understand the protective role of STAT3 and its relation to the Nrf2-signaling pathway, as well as the expression of antioxidant genes.

## Data availability statement

The data that support the findings of this study are available from the corresponding author upon reasonable request.

## Contributions

B.F. conceived the project. A.A. and B.F. conducted the experiments and collected the data. A.A and B.F. analyzed the data and wrote the manuscript. M.A. revised the manuscript. M.A. secured funding. All authors read and approved the final version of the manuscript.

---

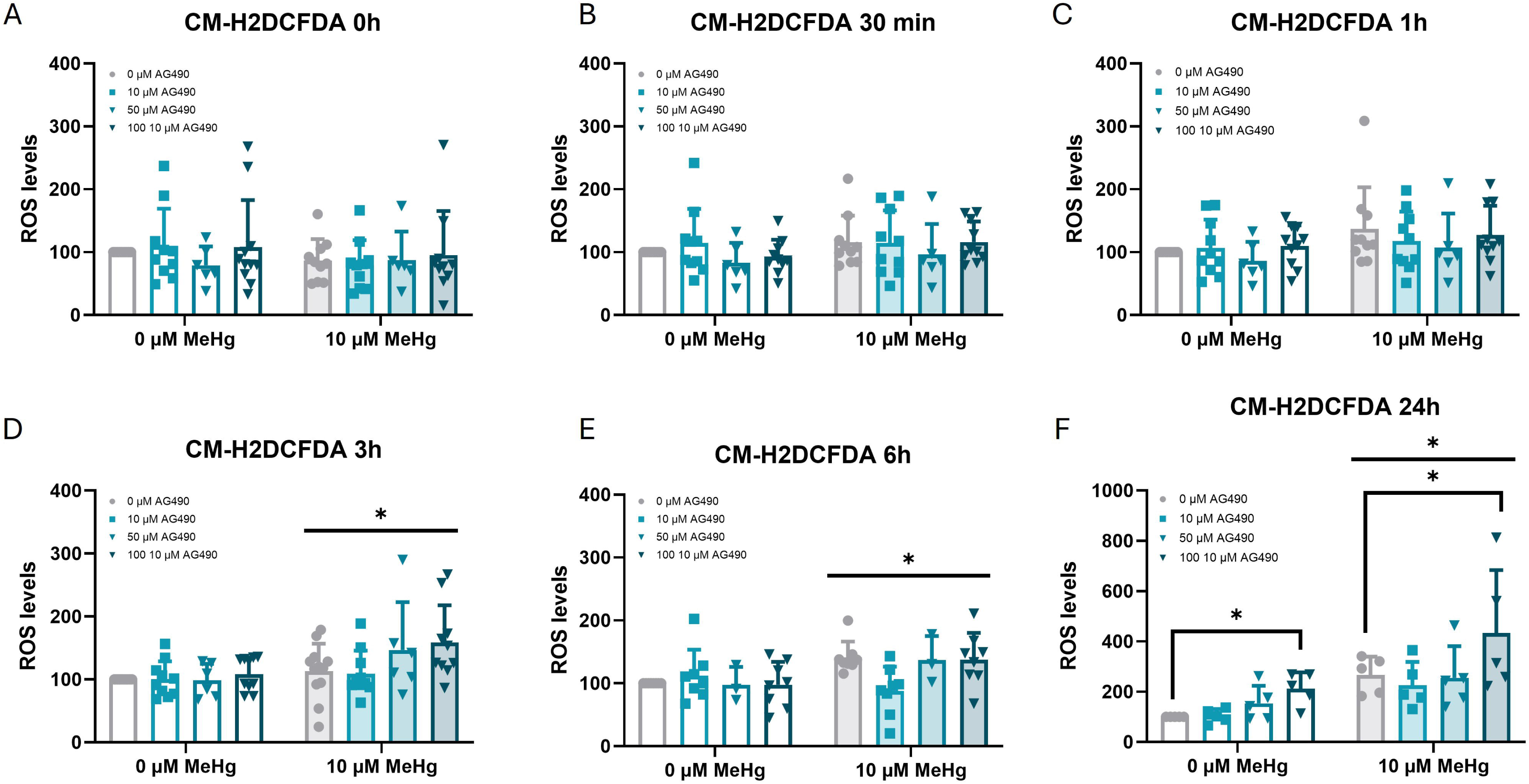

